# Semi-permeable capsules enable parallel cultivation and live microscopic observations of microbial eukaryotes

**DOI:** 10.64898/2026.04.02.716075

**Authors:** Marco Fantini, Nikolaj Brask, Sofia Paraskevopoulou, Humberto Itriago, Rustem Musaev, Julie Boisard, Karla I. Aguilera-Campos, Courtney W. Stairs

## Abstract

Semi-permeable capsules (SPCs) create enclosed porous microenvironments, diffusible to only small proteins and macromolecules. This presents a powerful tool for single cell observation, isolation, and sequencing. However, their range of use for sustaining viable microbial eukaryotes is largely unexplored. Single-cell eukaryotes are often understudied, with a wealth of unknown lifecycles, culturing methods and inter-microbial interactions, which are difficult to visualize. Here, we show that eukaryotes from eight different supergroups can be captured and propagated in SPCs. Encapsulation allowed observations of cell stages, motility and growth in a traceable and parallelized manner.

---

Microbial eukaryotes represent the majority of eukaryotic diversity, yet they remain among the least studied [1, 2], despite their importance for understanding the cell biology and evolution of eukaryotes [3]. Studying these organisms presents a variety of challenges as many single-celled eukaryotes exhibit multiple and complex life cycles that vary dramatically in motility, size, and morphology making taxonomic assignment tedious [4–6]. Moreover, establishing enrichment cultures from environmental sources is challenging; some protists are sensitive to predator-prey dynamics or cannot compete with blooms of fast-growing organisms. As a result, slow-growing or rare eukaryotes are outcompeted, leading to cultivation bias and an underrepresentation of potentially ecologically important lineages [7].

Semi-permeable capsules (SPCs) are microscale hydrogel compartments generated by microfluidics that form a shell around an inner aqueous core in which cells are confined. This porous shell allows free diffusion of small molecules (<340 kDa) and renders encapsulated cells chemically accessible while reducing aggregation, cross-contamination, and sample loss during downstream processing [8–10]. Previous studies have demonstrated the successful encapsulation of bacteria [10], mammalian cells [8], and viruses [11] but encapsulation of microbial eukaryotes has been limited [11].

To test whether SPCs can confine microbial eukaryotes, we encapsulated protists from ‘reference’ cultures and environmentally derived water samples (supplementary methods). Ten reference organisms (fig. 1) were selected to represent major eukaryotic lineages that span different nutritional modes (autotrophic, heterotrophic, and mixotrophic), movement types (amoeboid, euglenoid, flagellar, and non-motile), oxygen utilization, and salinity preference. Polyxenic cultures of breviates (*Pygsuia biforma*, and undescribed breviate *‘*Roskilde’) and *Euglena gracilis* were included to evaluate co-encapsulation of microbial eukaryotes and prokaryotes. This design allowed us to evaluate how cells from diverse phylogenetic, physiological, and ecological contexts responded to confinement in SPCs by recording their encapsulation efficiency, propagation, motility, division dynamics, and morphological variability.

**Figure 1.**
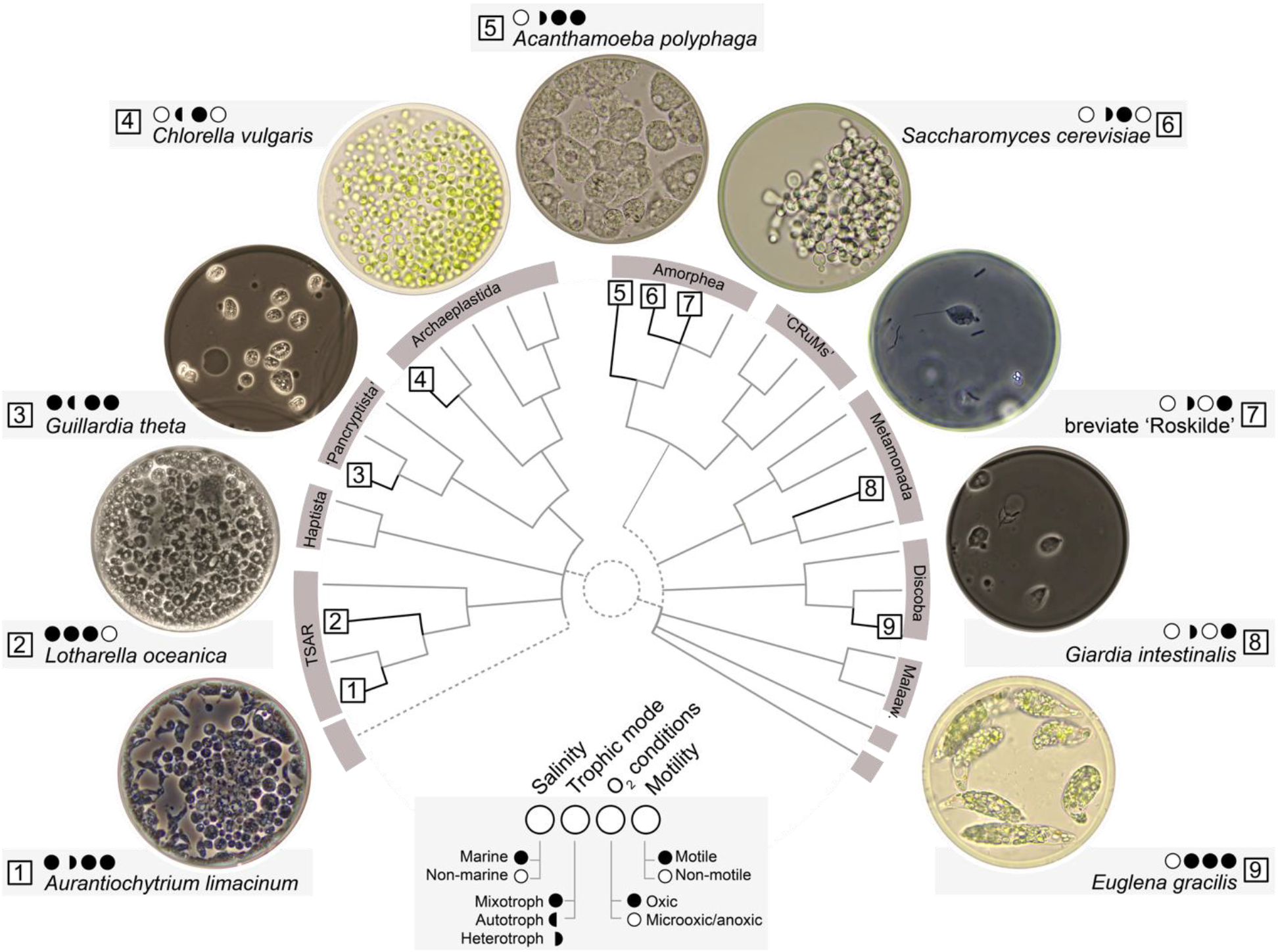
Eukaryotic tree of life, showcasing successful encapsulation of ‘reference’ organisms from various lineages and lifestyles labeled clockwise. For each organism the culture conditions (salinity, oxygen content) and generally accepted classification for trophic modes (mixotrophy, autotrophy, heterotrophy) and motility (motile, non-motile) are shown with closed and open circles according to the legend. Taxonomic relationships are adapted from Burki et al. and include reference organisms from TSAR (Stramenopila: *Aurantiochytrium limacinum*; Rhizaria: *Lotharella oceanica*), Cryptista (*Guillardia theta)*, Archaeplastida (*Chlorella vulgaris)*, Amorphea (Obazoa: undescribed breviate ‘Roskilde’, *Saccharomyces cerevisiae*; and Amoebozoa: *Acanthamoeba polyphaga*), Discoba (*Euglena gracilis)* and Metamonada *(Giardia intestinalis)*. Major eukaryotic groups are labeled grey on the tips [18].

We observed that most reference organisms survived encapsulation (9/10) and could divide (8/10). Motility was observed in all organisms with a motile vegetative stage despite confinement in the dense core of the capsule (5/5, ‘Roskilde’, *Acanthamoeba polyphaga, Euglena gracilis, Guillardia theta and Giardia intestinalis*, fig. 2, videos SV1-SV5). *G. theta* did not maintain its characteristic perpetual rotary swimming pattern and instead displayed sporadic rapid movements within the capsule (video SV4). Motility was observed in organisms with a motile zoospore stage (1/2) like *Lotharella oceanica* (video SV6) but not *Aurantiochytrium limacinum*. Overall, by limiting the dispersal of motile organisms, capsule confinement simplified microscopic examination over longer timepoints. Cell division processes were also observed inside capsules; for example, newborn *Chlorella vulgaris* cells can be seen dividing inside the initial mother cell wall (fig. 2C, video SV7) and similarly we observed cytokinesis in *E. gracilis* (video SV8) and *G. intestinalis* (fig. 2E). Morphological variability could be seen with *A. polyphaga*, where crowding of the SPC induced apparent encystation. Following capsule dissolution the *A. polyphaga* cells transition to an amoeboid state after 20 minutes (fig. 2D, video SV9).

**Figure 2.**
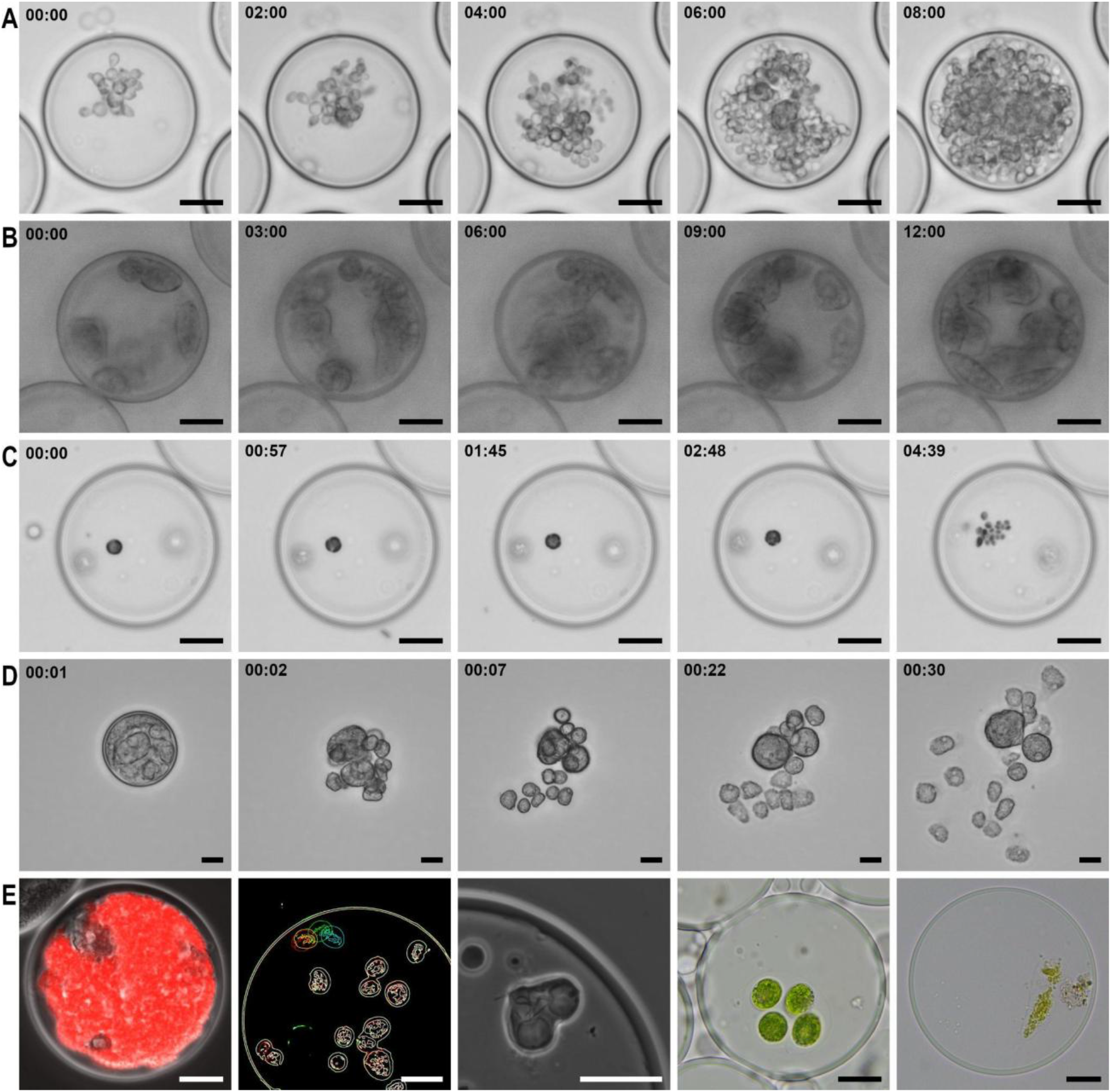
Time course images of microbial eukaryotes in semi-permeable capsules. A) *S. cerevisiae*, B) *A. polyphaga*, C) *C. vulgaris* growth in single capsules following encapsulation and transfer to original medium. D) Time series of *A. polyphaga* cysts confined in an SPC reverting to trophozoites upon capsule dissolution. E) Left to right: Fluorescent micrograph of *A. polyphaga* outcompeted by the co-encapsulated mScarlet-I tagged *E*.*coli* after 1 d of co-culturing; Movement of *G. theta* in capsule, color represents different timepoints; *Giardia intestinalis* trophozoite undergoing cytokinesis within an SPC, showing heart-shaped cell morphology, duplicated flagella and duplicated ventral discs; encapsulated microbial eukaryotes from environmental samples from a bog near Lake Tjörnarp (Prästabonnaskogen Nature Reserve) and from an urban pond in Lund, Sweden. Scale bar: 20 μm.

To understand the factors that influence cell viability within the capsule, we examined the effect of media osmolarity, oxygen dependency, feeding requirements, mixed cultures, and overcrowding on survival and proliferation. Of note, species that possess protective cellular structures correlated with viability in SPCs. For example, *A. limacinum, Saccharomyces cerevisiae*, and *L. oceanica* have a protective cell covering [4, 5, 12], *E. gracilis* has a pellicle, *Acanthamoeba* tropozoites carry contractile vacuoles and the cysts exhibit double walls, all of which provide support [13, 14]. To evaluate the effect of osmotic stress experienced by the cells, we examined cell viability of species derived from marine and non-marine habitats. Overall the non-marine eukaryotes, like *S. cerevisiae, A. polyphaga*, and *C. vulgaris*, showcased robust growth following encapsulation (fig. 2ABC) and generally resulted in higher survival following encapsulation compared to marine counterparts, *e*.*g. L. oceanica* and *G. theta*. Similarly, the marine breviate *P. biforma* did not survive encapsulation while the freshwater ‘Roskilde’, survived initial encapsulation but failed to replicate (fig. 1). Notably, the marine stramenopile *A. limacinum* deviated from this trend, displaying robust growth despite the salinity of its growth medium. This difference in viability of non-marine compared to marine species likely relates to osmolarity differences between their culture media and encapsulation solutions. Following encapsulation, we observed motility and cell division of the anaerobe *G. intestinalis* (fig. 2E, video SV5) suggesting that oxygen exposure during encapsulation (<1 h) was not a major source of cell damage for the protist. The presence of dithiothreitol (a reducing agent) in the encapsulation solutions likely further reduced such damage.

Unlike physical confinement strategies such as oil droplets, SPCs do not impose nutrient limitations intrinsic to droplet interiors, where nutrients are restricted to those initially present unless advanced microfluidic designs or micromanipulation are employed [15, 16]. Growth is, however, limited by capsule volume and by the capsule mechanical properties. Rapidly proliferating organisms in nutrient-rich conditions, such as *A. polyphaga* and *S. cerevisiae*, quickly reached overcrowding (videos SV2, SV10). Continued biomass accumulation generated internal pressure that exceeded capsule elasticity, resulting in rupture and loss of confinement (video SV10).

Heterotrophic organisms and mixed populations introduced additional constraints related to feeding and population balance. Organisms dependent on microbial prey, including the polyxenic ‘Roskilde’, were difficult to maintain within capsules. In such mixed populations, fast-growing prokaryotes frequently dominated capsule volume (supplementary methods, fig. 2E, S1, video SV11).

We encapsulated cells filtered from environmentally-sourced pond and bog water to assess whether SPCs could capture eukaryotes from natural communities (supplementary methods). In both cases, SPCs captured morphologically distinct eukaryotic cells (fig. S2, S3). If SPCs can be used to confine diverse natural communities, micromanipulation like single capsule picking could be used to selectively recover individual capsules containing organisms of interest. As a proof of concept single capsules of *A. polyphaga* and *E. gracilis* were handpicked using pulled capillaries and used to establish new cultures, demonstrating that encapsulation does not compromise downstream recovery (supplementary methods, videos SV9, SV12). This approach could facilitate the isolation of target species from mixed communities before predation or competitive overgrowth occurs, enabling the establishment of axenic or mono-eukaryotic cultures (video SV13). As SPCs are easy to pipette between wells in a microtitre plate without need of a micromanipulator, it allows users to perform screening of different media to monitor growth in-realtime with a standard inverted microscope.

SPCs offer a framework for permitting growth inside a capsule prior to downstream manipulation or omic analyses. This provides a new way to increase the quantity of sample material destined for single-cell studies. In addition, SPCs could provide an approach to isolate organisms that are challenging to single-cell ‘pick’ such as those that form chains, aggregates on debris, or transient multicellular networks [17]. SPCs address these limitations by enforcing physical separation and by generating uniform, isolated units, thereby extending single-cell omics approaches to non-model, debris-laden, and morphologically complex eukaryotes that are otherwise inaccessible to standard workflows.

Physical isolation of intact eukaryotic cells confines nucleic acids associated with the retained organism inside the capsule. This way SPCs provide a platform for high-throughput screening of microbial eukaryotes and their associated genetic elements, including mitochondria, plasmids, symbionts, microbiome, and viruses, extending previous applications of SPCs for studying microbes in mixed populations [11].

## Acknowledgments

We would like to acknowledge the support of John Archibald, Charlie Cornwallis, Elisabeth Gauger, Fiona Henriquez-Mui, Ondřej Pomahač, Ivan Čepička, Alastair G.B. Simpson and Staffan Svärd for providing cell cultures for encapsulation.

This project was supported by funds from the Swedish Research Council Swedish Research Council (Vetenskapsrådet starting grant 2020-05071 to C.W.S.), the European Research Council (ERC) under the European Union’s Horizon 2020 research and innovation programme (grant agreement ERC Starting grant 101078476 to C.W.S.), Human Frontiers in Research Programme (RGEC29/2024 to C.W.S.)

