## Supplementary information for "Semi-permeable capsules enable parallel cultivation and live microscopic observations of microbial eukaryotes"

The supplementary information includes the following:

**Supplementary Methods**

**Supplementary Figures S1-S3**

**Supplementary Videos SV1-SV13**

### Supplementary Methods

<u>Cell cultures:</u> *Chlorella vulgaris* strain Kgh101 was sourced from Charlie Cornwallis's culture collection (Lund University, Sweden); *Euglena gracilis* was provided by Elisabeth Gauger (Lund University, Sweden); *Giardia intestinalis* was provided by Staffan Svärd (Uppsala University, Sweden); *Acanthamoeba polyphaga* was provided by Fiona Henriquez-Mui (University of the West of Scotland, UK);  undescribed breviate Roskilde was provided by Ondřej Pomahač and Ivan Čepička (Charles University, Czechia); *Aurantiochytrium limacinum* (ATCC MYA-1381), *Lotharella oceanica* (CCMP622) and *Guillardia theta* (CCMP2712) were part of John M. Archibald culture collection and are available from the National Center for Marine Algae and Microbiota culture collection (NCMA, Canada) or from the American Type Culture Collection (ATCC, USA); *Pygsuia biforma* was provided by Alastair G.B. Simpson (Dalhousie University, Canada).

<u>Growth Media:</u>

Reference organisms were cultured in: *Aurantiochytrium limacinum*: modified 790 By+ (1.0 g/L yeast extract, 1.0 g/L peptone, and 5.0 g/L D-glucose dissolved in 33 g/L of

Instant Ocean), *Chlorella vulgaris*: Bold's Basal Medium (BBM) [1], *Acanthamoeba polyphaga*: PYG (7.5 g/L Bacto peptone, 7.5 g/L yeast extract, 15 g/L D-Glucose 0.15 g/L, $KH_2PO_4$ anhydrous 1.3 g/L, pH 6.7), *Euglena gracilis*: Pea infusion medium (40 green peas/L, boiled for 5 min, crushed, filter sterilized), *Saccharomyces cerevisiae*: YPD (10 g/L Yeast Extract, 20 g/L Bacto Peptone, 20 g/L D-glucose), undescribed breviate Roskilde: ATCC Medium 1773 [2], *Lotharella oceanica, Guillardia theta*: f/2 medium [3], *Giardia intestinalis*: TYDK medium (30 g/L peptone, 10 g/L glucose, 2 g/L NaCl, 0.2 g/L L-ascorbic acid, 1 g/L $K_2HPO_4$, 0.6 g/L $KH_2PO_4$, 2 g/L L-cysteine, 22 mg/L ferric ammonium citrate, 125 mg/L bile) supplemented with 10% heat-inactivated FBS, *Pygsuia biforma*: modified MYB (0.1 g/L yeast extract, 0.1 g/L maltose dissolved in 33 g/L of Instant Ocean).

<u>Encapsulation and Imaging:</u>

Upon reaching densities between 1 x $10^6$ - 3 x $10^7$ cells/mL, cultures were subjected to centrifugation (500-3000 x g), for 5 - 15 min. Final cell densities before encapsulation were as follows (millions cells/mL): *Aurantiochytrium limacinum*: 36.7, *Chlorella vulgaris*: 15.7, *Acanthamoeba polyphaga*: 6.4, *Euglena gracilis*: 7.6, *Saccharomyces cerevisiae*: 23.9, undescribed breviate Roskilde: 1.4, *Lotharella oceanica*: 2.1, *Guillardia theta*: 1.6, *Pygsuia biforma*: 5.9. *Giardia intestinalis* was not counted prior to encapsulation because of biosafety limitations. The culture suspension was processed using an SPC Innovator Kit (60 µm, Atrandi Biosciences, Lithuania) and injected into the Onyx encapsulation device (Atrandi Biosciences, Lithuania). Two distinct sample preparation protocols were used for encapsulation: for general use we followed the company recommended protocol (50 µL core solution 2x, 12.5 µL photoinitiator, 1 µL DTT, and phosphate buffer saline (PBS) 1x and culture sample to reach the desired occupancy λ of 0.1-0.2); for marine organisms we designed a high-salinity variation (40 µL core solution 2x, 10 µL photoinitiator, 1 µL DTT, 44 µL concentrated filtered Instant ocean 1.5x (49.5 g/L), 5 µL concentrated sample, to reach a final salinity of ~71% sea level). Capsule formation and emulsion breaking were performed according to the supplier instructions. 100-200µL out of the ~500µL obtained capsules were diluted to 5 mL and moved to tubes (*Giardia intestinalis*) or culture flasks (TC-treated T-25 for *Pygsuia biforma* and undescribed breviate Roskilde; T-25 with vented cap for the remaining organisms) and 100 µL was used for initial microscopy, via microscopy slides with raised coverslip using office tape. Capsules were transferred to adapted microscope slides for a second round of imaging once growth had been observed, typically after one to three d for most cell lines, 8 d for *Guillardia* and 24 d for *Lotharella*. To recover cells from the capsule, capsule dissolution was induced with the release agent included in the SPC innovator kit (1% added directly into the culture medium).

Microscopy:

Capsules were visualized in phase and brightfield on an Olympus CX43 or Zeiss Axio Imager .v2 microscope. 18-20 h timelapses were taken of *A. polyphaga*, *C. vulgaris* and *S. cerevisiae* on a Zeiss Axio observer .z1 inverted microscope. Similarly, recordings of movement in the capsule were taken for *E. gracilis*, undescribed breviate 'Roskilde', *G. theta*, *L. oceanica, G. intestinalis* in either the Olympus or Zeiss Microscopes.  Capsule dissolution was recorded for *A. castellani*, *E. gracilis*, *C. vulgaris* and *S. cerevisiae* on the Zeiss Axio observer inverted microscope. A single capsule of *A. castellani* and *E. gracilis* was picked and moved to fresh medium previous capsule dissolution. A *G. theta* timelapse microscopy (phase contrast, video SV4) was processed with Fiji/imageJ v1.54p to visualize movement inside the capsule. In order, background subtraction (ball radius 50 px), enhanced contrast, intensity threshold (top 6%) and edge detection was performed on selected frames which were then merged in a single image. Imaging of *G.intestinalis* was performed in a Nunclon Delta Flat-Sided Tube (VWR). The curvature of the opposite side of the vial produced directional shadows during brightfield microscopy.

Environmental sample collection and processing:

Environmental samples were collected from freshwater habitats in southern Sweden, from a bog in the Prästabonnaskogen Nature Reserve near Lake Tjörnarp, Sweden (55.99740° N, 13.62391° E), September 29th 2025, 10 AM. At the time of sampling, light intensity was 19-122 lux, ambient temperature was 13.5°C, and water temperature was 11.0°C. Sampling was performed at a depth of 9 cm. A second sample was collected from an urban pond in the man-made meadow outside Ecologihuset, Lund, Sweden (55.71378° N, 13.20683° E), October 21th 2025, 10 AM. At the time of sampling, light intensity was 14 lux, ambient temperature was 12.0°C, and water temperature was 8.0°C. Sampling was performed at a depth of 8 cm. At each site, 20 L of water was collected. Samples were first passed through a 74 µm mesh sieve (20 cm diameter) four times to remove large debris and then through a 40 µm cell strainer. The material was subsequently concentrated to approximately 15 mL using a Vivaflow 200 system fitted with a 0.2 µm membrane and encapsulated in SPCs with the standard protocol. For the urban pond sample, only 6 L of the collected water was processed by sieving prior to concentration with the Vivaflow system. After encapsulation the capsule were moved in modified artificial pond water (2 g/L $Ca(NO_3)_2 \cdot 4H_2O$, 1 g/L $K_2HPO_4$ anhydrous, 1 g/L $Na_2SiO_3 \cdot 9H_2O$, 5mL f/2 micronutrient solution [3]).

#### Acanthamoeba and mock prokaryotic community

*Pseudomonas aeruginosa* and *Escherichia coli* were grown overnight in PYG medium, diluted 1:20 and 1:10 respectively in fresh PYG, and incubated for 1 h at 37°C. Bacterial density was estimated by optical density and Bürker chamber counts. *Acanthamoeba polyphaga* was grown for 7 d at 16°C and counted in a Bürker chamber. Bacteria were diluted in PBS (Phosphate Buffered Saline, pH 7.4) and amoebae concentrated by low-speed centrifugation (800 x g, 5 min). Cell suspensions were diluted with the encapsulation solution to a concentration of $8 \times 10^6$ cells/mL for each bacterial species and $5 \times 10^7$ cells/mL for *A. polyphaga* (4:4:25 ratio). The mixture was immediately encapsulated, and batches of 20 µL of packed SPCs were transferred to 12-well plates containing 1 mL PBS with 1% PYG. Growth was monitored daily by microscopy for 3 d. *Pseudomonas aeruginosa* was provided by Elisabeth Gauger (Lund University, Sweden); and *Escherichia coli* is a standard laboratory strain (DH5α, Thermofisher).

**Supplementary Figures**

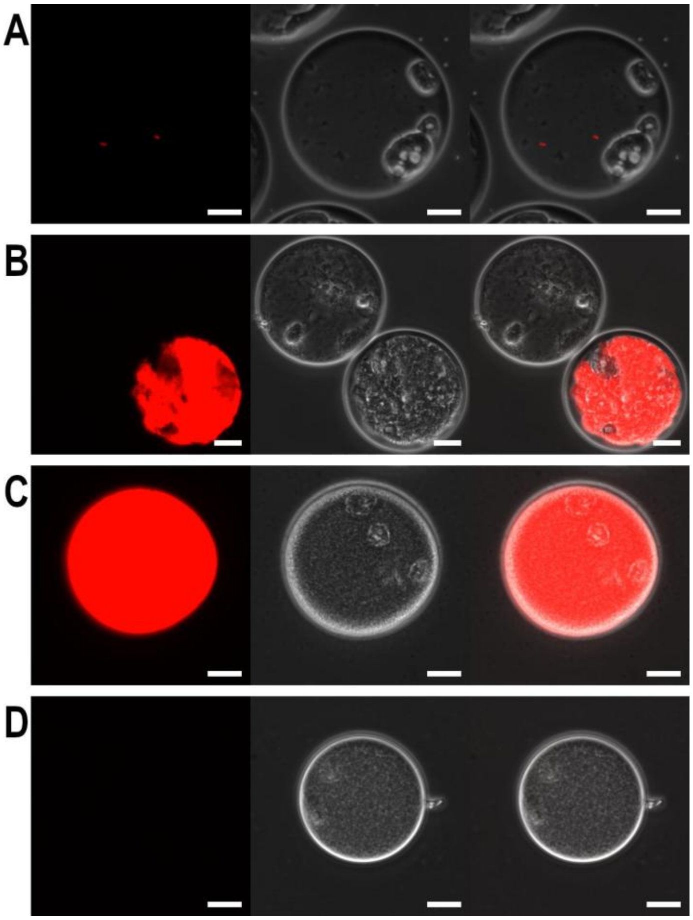


**Supplementary Figure S1.** Timecourse of bacterial overgrowth in SPCs containing *Acanthamoeba polyphaga* with *Escherichia coli* and *Pseudomonas aeruginosa (25:4:4)*. *E. coli* tagged with mScarlet-I shows red fluorescence, *P. aeruginosa* is untagged and shows no fluorescence. A) Day 0, single bacterial cells within the capsule. C) Day 1, onset of bacterial overgrowth for both *E. coli* and *P. aeruginosa*. C-D) Day 3, extensive bacterial overgrowth overwhelming *A. polyphaga*, with *E. coli* in (C) and *P. aeruginosa* in (D). Left to right: fluorescence (Ex. 586/20, Di. 605, Em. 647/57), phase contrast and merged images, scale bar 20 μm.

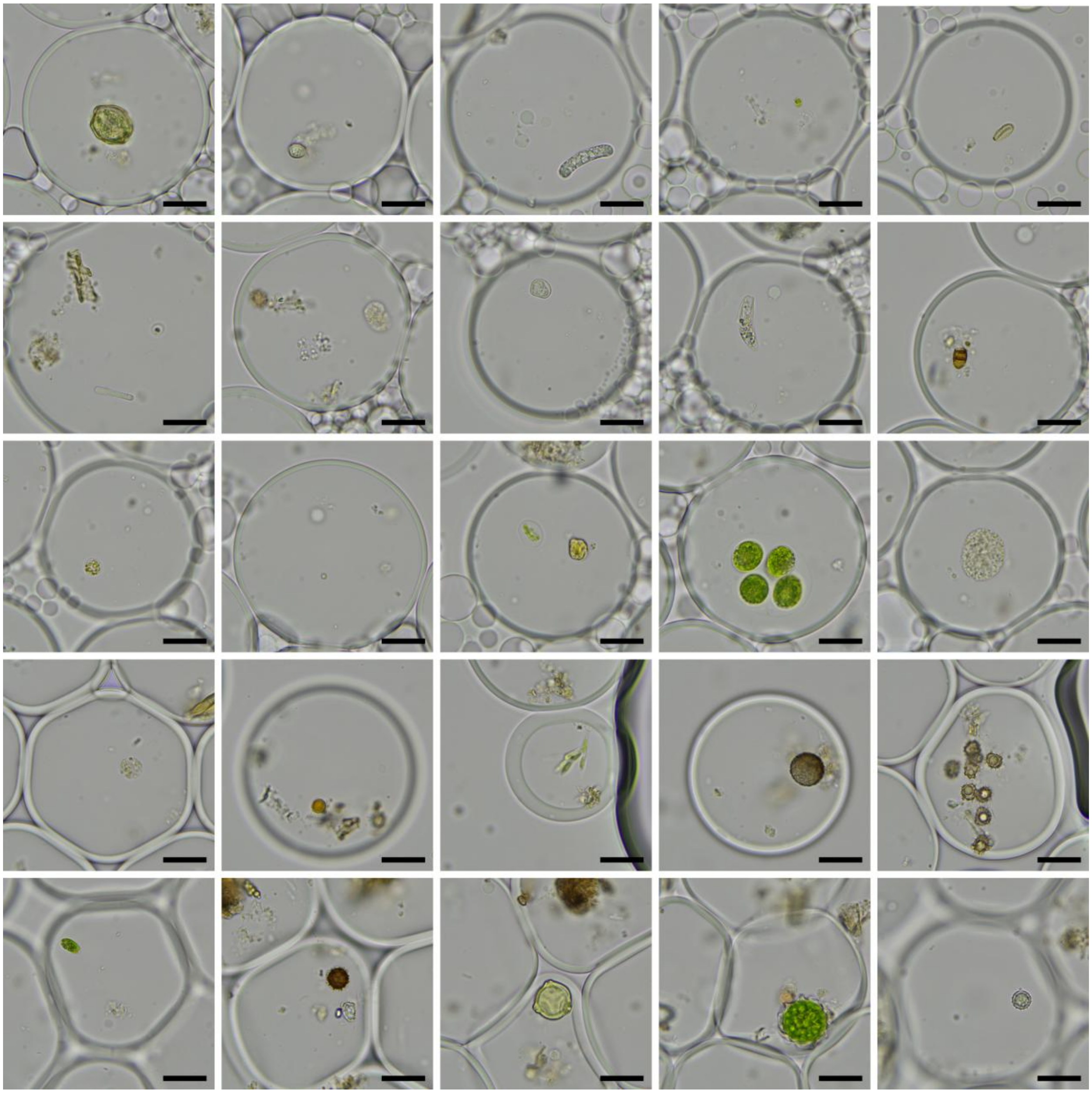

**Supplementary Figure S2.** Eukaryotic microorganisms from a bog near Lake Tjörnarp (Sweden) encapsulated in SPCs. Representative brightfield micrographs, scale bar 20 µm.

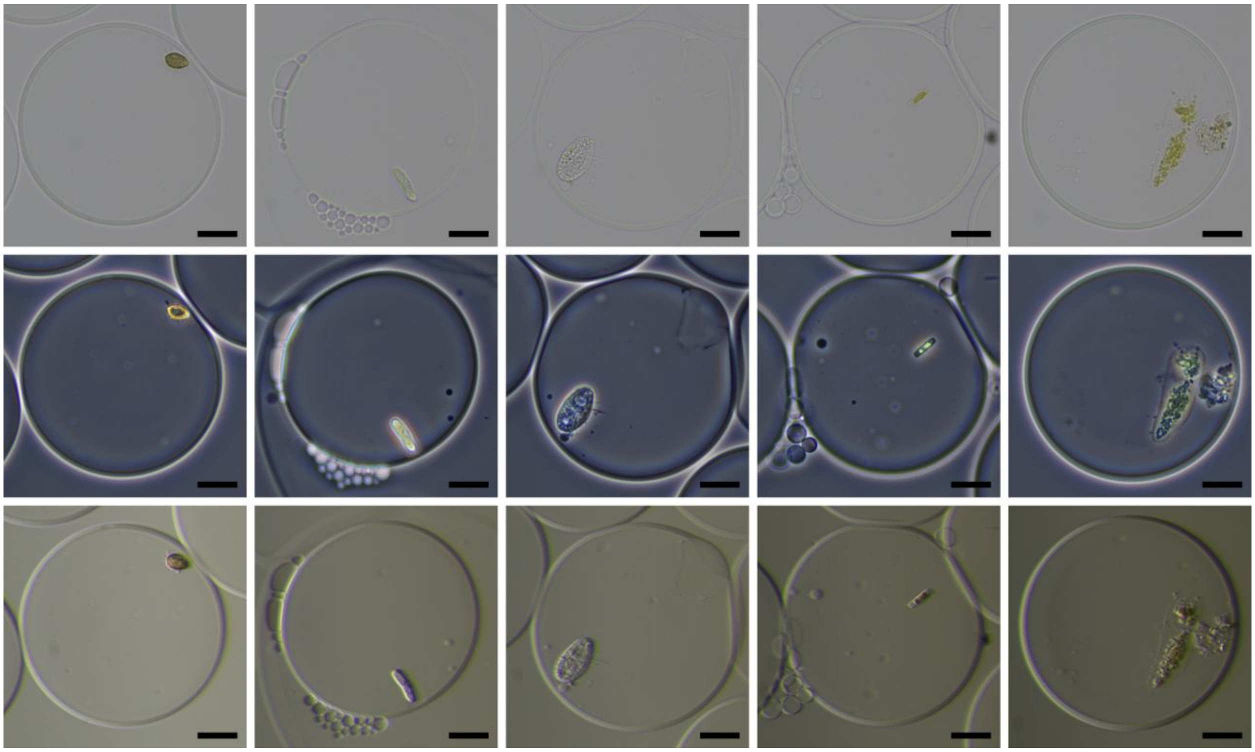

**Supplementary Figure S3.** Eukaryotic microorganisms from an urban pond (Lund, Sweden) encapsulated in SPCs. Representative brightfield, phase contrast and oblique illumination micrographs, scale bar 20 µm.

**Supplementary Video SV1.** Undescribed breviate Roskilde encapsulated with some bacteria from its cohort in SPCs. Phase contrast microscopy, scale bar 20μm

**Supplementary Video SV2.** Timelapse of *A. polyphaga* crawling and dividing inside a SPC. *A. polyphaga* trophozoites show adhesion to the capsule shell. Scale bar 20μm.

**Supplementary Video SV3.** *E. gracilis* showing both flagellar and peristaltic movement inside a SPC. Bacterial growth was controlled with 1% PenStrep. Scale bar 20μm.

**Supplementary Video SV4.** *G. theta* inside an SPC showing sporadic rapid movements within the capsule rather than its characteristic rotary swimming pattern. Phase contrast microscopy, scale bar 10μm.

**Supplementary Video SV5.** Timelapse of *G. intestinalis* inside SPCs, showing both attached and free-swimming trophozoites. Brightfield microscopy, scale bar 50μm.

**Supplementary Video SV6.** *L. oceanica* inside an SPC, with a motile zoospore moving around the capsule interior. Scale bar 20μm.

**Supplementary Video SV7.** Timelapse of *C. vulgaris* growing and dividing inside SPCs. Scale bar: 20μm

**Supplementary Video SV8.** Capsule filled with E. *gracilis* in the process of cytokinesis. Scale bar: 20μm

**Supplementary Video SV9.** Timelapse of *A. polyphaga* cysts confined in an SPC reverting to trophozoites upon capsule dissolution. The capsule was isolated with manual micropipetting prior to capsule dissolution. Scale bar: 50μm.

**Supplementary Video SV10.** Timelapse of *S. cerevisiae* dividing inside a SPC. *S. cerevisiae* budding inside the capsule form large chains of loosely attached cells. When the pressure of the growing yeast biomass overcomes the elasticity limit of the capsule, a rupture occurs and cells are dispersed in the surrounding medium. Scale bar 20μm.

**Supplementary Video SV11.** Timelapse of a polyxenic culture of *E. gracilis* encapsulated in SPCs showing fast growing bacteria rapidly filling the capsule volume. Scale bar 20μm.

**Supplementary Video SV12.** *E. gracilis* being released in the media after single capsule isolation and dissolution. Scale bar: 50 µm.

**Supplementary Video SV13.** Polixenic *E. gracilis* culture after confinement in SPCs for 1 day without antibiotics. The capsule shown contains *E. gracilis* cells without the dense bacterial growth seen in surrounding capsules, indicating the potential for selective recovery of capsules containing only the target eukaryote. Scale bar 20µm.


1. Wong Y, Ho Y, Ho K, et al. Growth Medium Screening for Chlorella vulgaris Growth and Lipid Production. *JAMB* 2017;**6**. https://doi.org/10.15406/jamb.2017.06.00143
2. ATCC. ATCC Medium: 1773 Hexamita Medium. https://www.atcc.org/-/media/product-assets/documents/microbial-media-formulations/1/7/7/3/atcc-medium-1773.pdf. .
3. Guillard RRL. Culture of Phytoplankton for Feeding Marine Invertebrates. In: Smith WL, Chanley MH (eds), Culture of Marine Invertebrate Animals: Proceedings — 1st Conference on Culture of Marine Invertebrate Animals Greenport. Boston, MA: Springer US, 1975, 29–60.
